# The use of hyperimmune chicken reference sera is not appropriate for the validation of influenza pseudotype neutralization assays

**DOI:** 10.1101/180026

**Authors:** Francesca Ferrara, Eleonora Molesti, Simon Scott, Giovanni Cattoli, Nigel Temperton

## Abstract

Pseudotype particle neutralization (pp-NT) is a next-generation serological assay employed for the sensitive study of influenza antibody responses, especially haemagglutinin stalk-directed antibodies. However, to date a validation of this assay has not been performed, and this limits its use to primarily research laboratories. To identify possible serological standards to be used in optimization and validation of the pp-NT, we have evaluated the cross-reactivity of hyperimmune chicken reference antisera in this assay. Our findings show that the cross-reactivity detected by the pp-NT assay is only in part explained by phylogenetic relationships and protein homology between the HA subtypes analysed; further studies are necessary to understand the origin of the cross-reactivity detected, and reference standards with higher specificity should be evaluated or generated *de novo* for future use in pp-NT.

## Introduction

Serological methods, such as single radial haemolysis (SRH), haemagglutination inhibition (HI) and microneutralization (MN), are cost-effective and widely-used methodologies to monitor the circulation and the prevalence of influenza viruses, and are also employed in vaccine immunogenicity studies in human and animal populations [1]. However, these assays are subject to numerous shortcomings, especially related to variability of the reagents, their standardization between laboratories, and their ability to detect haemagglutinin (HA) stalk-directed antibodies.

Recently, the use of replication-deficient viruses in neutralization assays is being widely evaluated as a potential alternative to MN assays. For example, HA pseudotype particles (pp) harbouring the HA embedded in the viral envelope and having a lentiviral vector as backbone, have been used effectively for this purpose [2,3]. The pseudotype particle neutralization (pp-NT) assay appears to be more sensitive than other functional assays to detect the antibody response directed against the HA stalk [4]. Consequently, pp could be used to effectively study heterosubtypic antibody responses directed against the HA stalk region. pp-NT assays have been shown to correlate strongly with other classical serological assays [5-8]. Since pp are replication-defective, they offer a safe alternative to wild-type virus methods (that require Biosafety Level 3 containment), and the detection of antibody responses is not influenced by the variability of blood-based reagents as observed in other assays (i.e. red blood cells in HI). Unfortunately, the validation of pp-NT, which will permit its more extensive use in surveillance, and in pre-clinical and clinical studies, has not been undertaken to date. Validation of pp-NT will allow certifying its use in product release and stability control (if, in future, HA stalk-directed monoclonal antibodies will be licensed) and in evaluating antibody responses elicited by current and ‘ next-generation universal’ influenza vaccines.

Essential for the optimization and the validation of an assay is the availability of appropriate reference materials. Appropriate reference standards are especially useful when the specificity, sensitivity, precision, and accuracy of an assay are evaluated for the first time [9] but are also essential when other assay parameters, such as dilution range or calibration curves, are established [10]. Furthermore they have an essential role to monitor assay stability and consistency over different analytical sessions (e.g. days) [11]. Reference materials are also useful when multisite validation of an assay is performed. For example the use of reference standard sera has been shown to be extremely useful to increase consistency between laboratories using HI and SRH [12].

Chicken reference antisera against all the HA subtypes are commonly generated and used for influenza virus typing in the HI assay [13,14], and thus should be investigated as possible controls and reference materials in pp-NT assays, which was the purpose of this study.

## Results and Discussion

We have evaluated the neutralization activity and cross-reactivity of chicken reference antisera against a panel of pp bearing HAs of a representative strain, where possible of avian origin, for each HA subtype. Unfortunately, H6 and H13 influenza pp were not used in this study since appropriate pp bearing the HA of these two subtypes was not available.

The neutralizing titres resulting after analysis with GraphPad Prism software were elaborated using the Microsoft Excel and statistical software R to generate a ‘ heat-map’ (Figure 1). Phylogenetic analysis was also used to highlight relationships between HA subtypes. The cross-reactivity heat map shows that influenza reference antisera are usually able to efficiently neutralize HA-matched pp, and minor variation in the neutralizing titres can be observed when the serum was generated against paired (same subtype but different strain) HAs. Additionally, cross-reactive responses can be detected not only when phylogenetic relationships are present between the HA of the pp tested and the HA used as antigens to generate the antisera, but also between HA and antisera that share less similarity (Figure 1).

**Figure 1:**
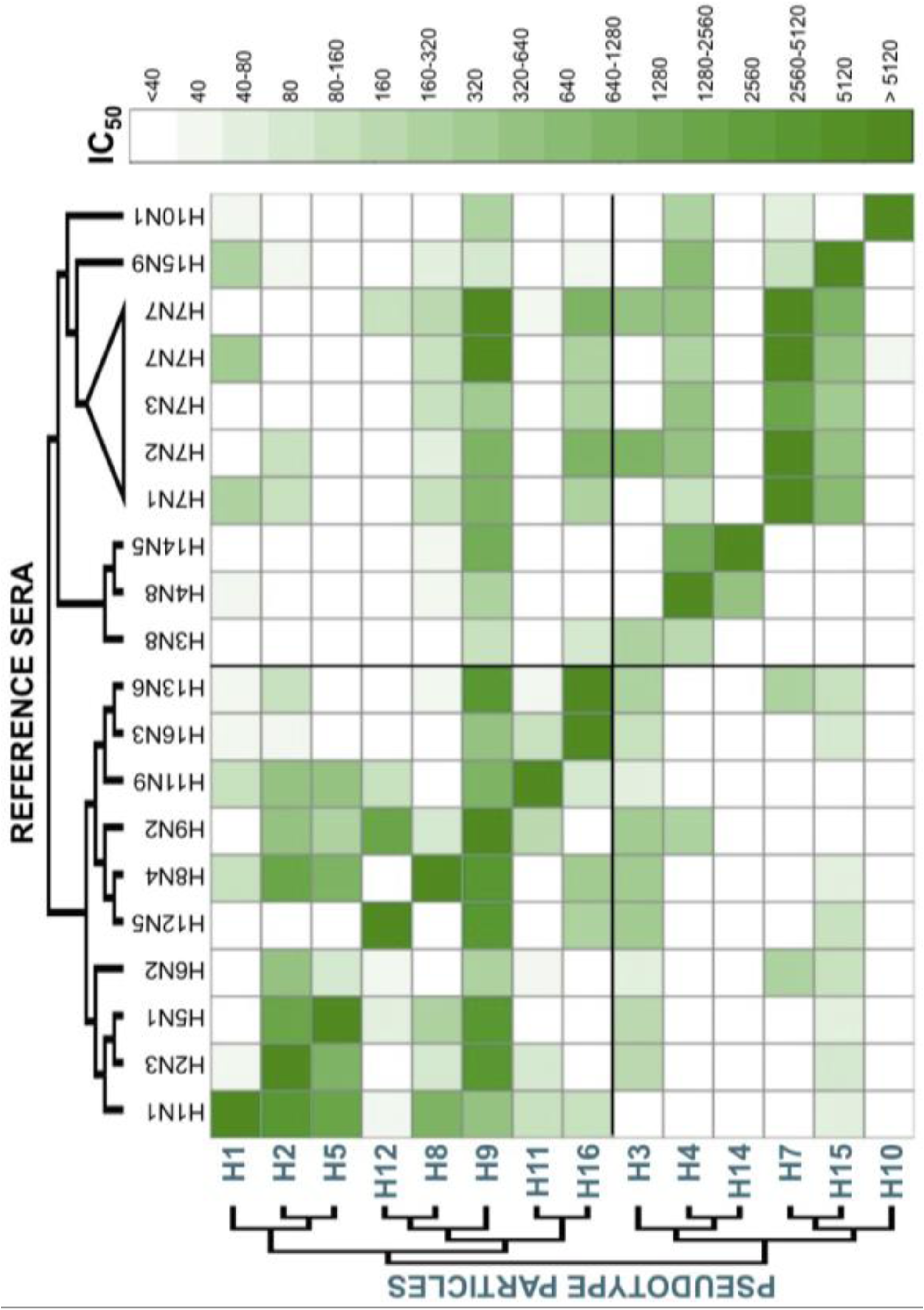
Cross-reactivity map of pp and reference sera based on IC_50_

Many studies have previously observed that chicken antisera generated using whole virus, in comparison to the ones generated through HA-expressing DNA vaccination or recombinant HA1 vaccination, present a lower specificity in HI and/or immunodot-blot assays: this is primarily due to the fact that antisera produced using whole virus also includes NA- and M2-directed antibodies [15,16]. However since a certain level of cross reactivity is observed also with DNA or recombinant protein vaccination, cross-reactive HA-directed antibodies are most probably involved [16].

The fact that reference antisera show high neutralizing responses and cross-reactivity between different strains/subtypes could be problematic not only for HI typing, but especially for pp-NT assays. In fact, in the dilution range analysed the pp-NT assay cannot discriminate between two pp. Preparation of the standard material through dilution of the commonly used reference standard, or the use of monoclonal antibody mixtures (with or without the presence of a serum matrix) showing high specificity could be more effective approaches to establish reference materials to be used to validate, standardise, and control the pp-NT assay. To better understand which factors are underlining the cross-reactivity observed we have evaluated the percentage amino acid identity between the HAs present on the pp, and the HAs used for antisera generation. A percentage-identity heat map was then designed (Figure 2). This map shows a similar pattern to the cross-reactivity map (Figure 1). Since it is difficult to highlight all the differences and similarities by eye, statistical analysis was performed to see whether any concordance or association between the maps was present. Kendall τ test shows that there is low association between percentage identity and neutralization titres (τ = 0.269, p ≤ 2.22 10^-16^). This means that the percentage of amino acid identity is a good approach to evaluate the cross-reactivity response, but to understand and explain the cross-reactivity detected, other approaches should be used. Recently antigenic cartography has been used to evaluate the antigenic evolution/drift of different influenza viruses and to help vaccine strain selection [17,18]. This methodology could be used to analyse the data presented here, and should permit a representation of the antigenic interplays between different pp. In the absence of appropriate controls and presence of high cross-reactivity responses, it will be difficult to assess the specificity of the pp-NT assay. Other parameters should be evaluated to understand if factors unrelated with the sera antibody content could interfere with the pp-NT assay. For example the presence of virus-attachment inhibitors in the sera and serum treatments (e.g. heat-inactivation, pre-treatment with receptor-destroying enzymes) can be assessed to optimise pp-NT assay conditions and to reduce non-specific neutralization if present. Also the evaluation of possible haemolysis or other contaminants (e.g. lipids) of the serum samples is a factor that needs to be taken into consideration when the assay is optimised and validated [9]. To conclude, the results presented here show that the high sensitivity and the propensity of the pp-NT assay to detect cross-reactive responses does not permit the use of current chicken reference standard antisera as reference materials to validate the assay. Until more appropriate standards (e.g. monoclonal antibody mixtures) will be developed to further progress optimisation and validation of the pp-NT assay the routinely used reference standards should be used as positive neutralization controls only in experimental research settings.

**Figure 2:**
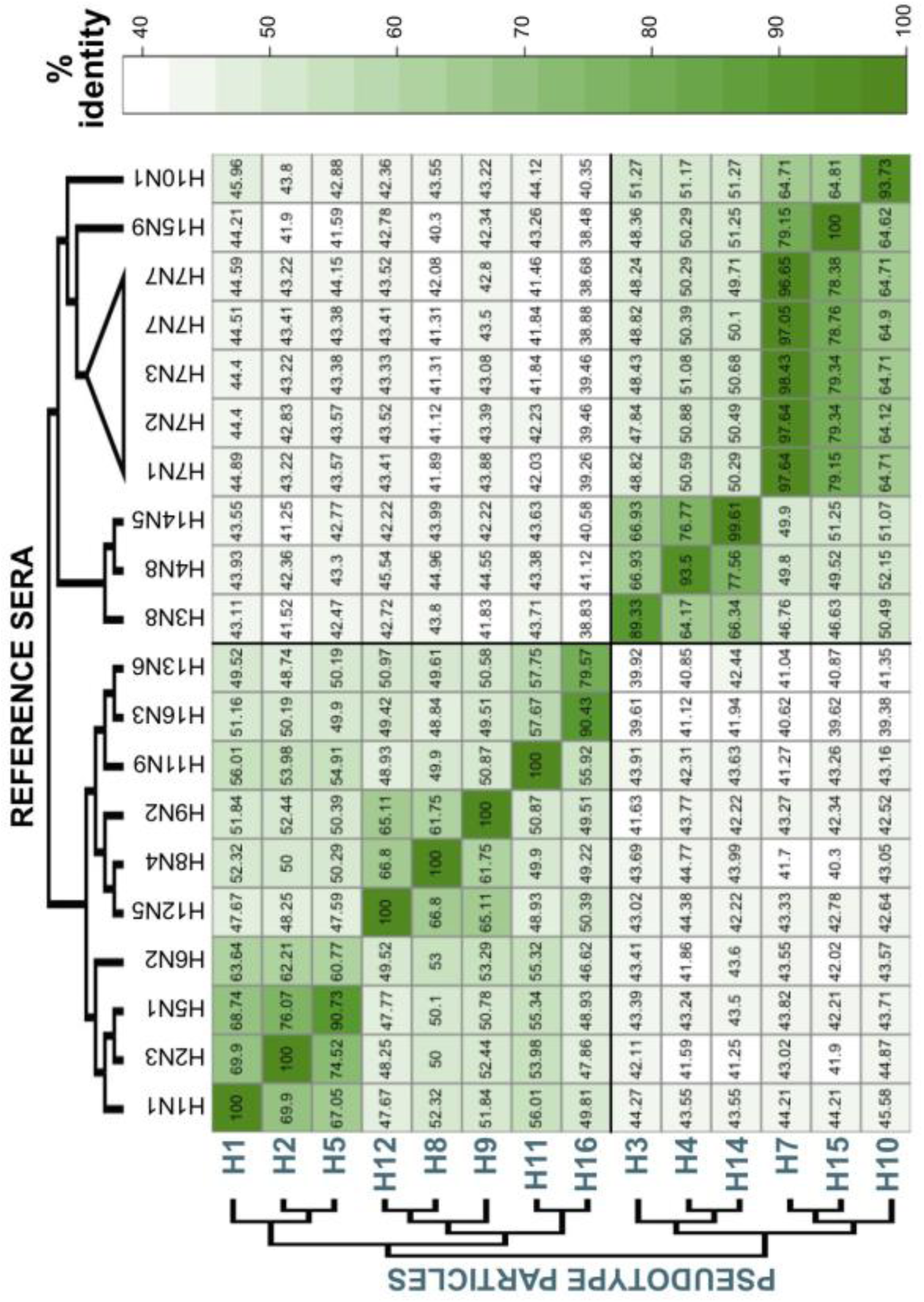
Cross-reactivity map of pp and reference sera based on percentage of amino acid identity

## Materials and Methods

### Reference sera

The OIE avian reference hyperimmune sera used for these studies and associated HI titres were provided by Dr. Giovanni Cattoli when he was at the Istituto Zooprofilattico delle Venezie, OIE, Legnaro, Padua, Italy and are reported in Table 1.

**Table 1:** OIE avian influenza reference antisera for HI assay, Agar Gel Immunodiffusion test, and Agar Gel Precipitation test

| Antigen strain name | Subtype | HA accession number | HI titre |
| --- | --- | --- | --- |
| **A/duck/Italy/1447/2005** | H1N1 | HF563054.1 | 1:512 |
| **A/duck/Germany/1215/1973** | H2N3 | CY014710.1 | 1:512 |
| **A/psittacine/Italy/2873/2000** | H3N8 | GQ247846.1* | 1:256 |
| **A/cockatoo/England/1972** | H4N8 | GQ247847.1* | 1:128 |
| **A/turkey/Canada/1965** | H6N2 | GQ247851.1* | 1:256 |
| **A/turkey/Ontario/6118/1968** | H8N4 | CY014659.1 | 1:512 |
| **A/mallard/Italy/3817-34/2005** | H9N2 | Not Applicable | 1:256 |
| **A/ostrich/South Africa/2001** | H10N1 | GQ247860.1* | 1:512 |
| **A/duck/Memphis/546/1974** | H11N9 | AB292779.1 | 1:1024 |
| **A/duck/Alberta/60/1976** | H12N5 | CY130078.1 | 1:128 |
| **A/gull/Maryland/704/1977** | H13N6 | D90308.1 | 1:1024 |
| **A/mallard/Gurjev/263/1982** | H14N5 | M35997 | 1:512 |
| **A/shearwater/Australia/2576/1979\*\*** | H15N9 | CY130102.1 | 1:2048 |
| **A/gull/Denmark/68110/2002** | H16N3 | GQ247872.1* | 1:256 |

\*Partial sequence
\*\*Also known as A/shearwater/West Australia/2576/1979

Reference avian sera against H5 and H7 influenza strains were provided by the Animal and Plant Health Agency (APHA, previously Animal Health and Veterinary Laboratories Agency) and are reported in Table 2.

**Table 2:** APHA avian influenza reference antisera

| Antigen strain name | Subtype | HA accession number | HI titre |
| --- | --- | --- | --- |
| A/chicken/Scotland/1959 | H5N1 | CY015081.1 | Not available |
| A/African starling/England/983/1979 | H7N1 | AF202232.1 | Not available |
| A/chicken/Wales/1306/2007 | H7N2 | EF675618.1 | Not available |
| A/chicken/England/4054/2006 | H7N3 | EF467826.1 | Not available |
| A/England/268/1996 | H7N7 | AF028020.1 | Not available |
| A/turkey/England/647/1977 | H7N7 | AF202247.1 | Not available |

### Pseudotype particles and pseudotype particle neutralization assays

Pseudotype particles were generated as described elsewhere [19] using HAs reported in Table 3.

**Table 3:**
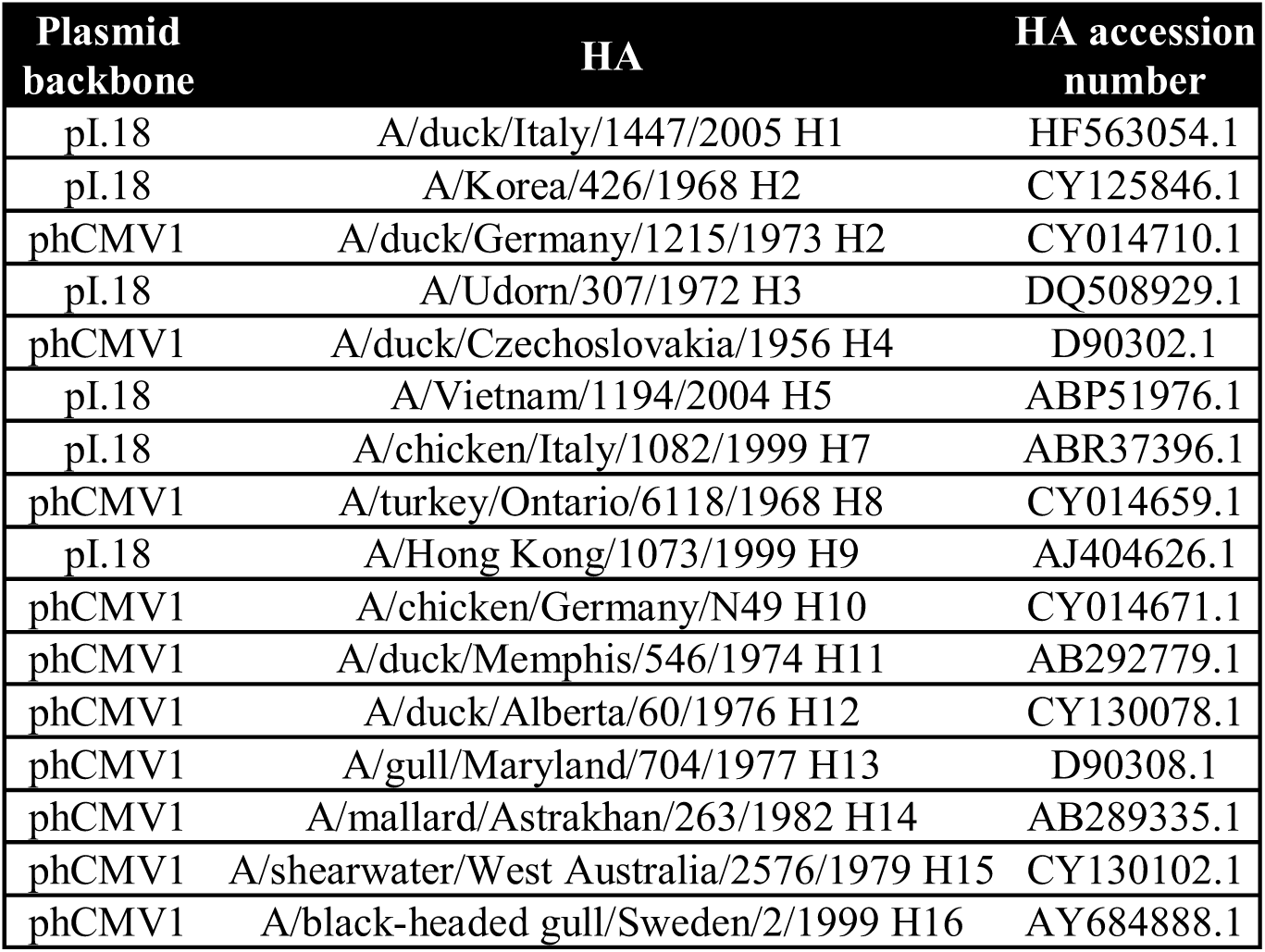
Haemagglutinin encoding plasmids used for pseudotype production

The pp-NT assays were performed as described elsewhere [19], using 5 μl of each serum sample (starting dilution 1:40) and using a pp input of 1×10_6_ RLU/well. IC_50_ neutralization titres were calculated using GraphPad Prism^®^ expressed as dilution factor; then for further statistical analysis they were categorised into 17 groups according to the dilution tested and as reported in Table 4.

**Table 4:** Category of IC_50_ used for statistics

| Group | IC<sub>50</sub> values | Dilution Factor |
| --- | --- | --- |
| 0 | <35 | <40 |
| 1 | 35-45 | 40 |
| 2 | 45-75 | 40-80 |
| 3 | 75-85 | 80 |
| 4 | 85-150 | 80-160 |
| 5 | 150-170 | 160 |
| 6 | 170-310 | 160-320 |
| 7 | 310-330 | 320 |
| 8 | 330-630 | 320-640 |
| 9 | 630-670 | 640 |
| 10 | 670-1270 | 640-1280 |
| 11 | 1270-1290 | 1280 |
| 12 | 1290-2550 | 1280-2560 |
| 13 | 2550-2570 | 2560 |
| 14 | 2570-5100 | 2560-5120 |
| 15 | 5100-5140 | 5120 |
| 16 | >5140 | >5120 |

A cross-reactivity map (pp versus reference antisera), completed using the neutralization groups for further statistical analysis, was designed in a Microsoft^®^ Excel 2011 spread sheet and then saved as a comma-separated value (csv) file.

### Bioinformatic analysis

Percentage identity between HA amino acid sequences of pp and reference sera antigens were calculated to check if cross-reactivity could be explained by overall sequence similarity. Amino acid sequences of the HA used in neutralization assays, and used to generate the reference antisera, were downloaded from the Influenza Virus Resource, the Influenza Research database and the Global Initiative on Sharing Avian Influenza Data (GISAID) Epi-Flu™ platform. The accession numbers of the HAs used in pp-NT assays are reported in Table 3; HA accession numbers of the reference antisera are reported in Tables 1 and 2. For A/mallard/Italy/3817-34/2005 (H9N2) the HA sequence was not available at the time of analysis and therefore the pp sequence was used as reference. Additionally, some of the HA sequences in the databases were incomplete, which complicates the analysis. To minimize this, it was decided to evaluate the percentage identity only for the amino acids that constitute the extracellular part of the HA (amino acids from 24 to 547 -H3 numbering), which were available for all HAs used. All sequences were aligned using MUSCLE algorithm [20] and Jalview software [21]. Subsequently the sequences were trimmed of their N-Terminal signal sequence, the transmembrane region, and the cytoplasmic tail. Percentage identities between amino acid sequences were calculated by pair-wise alignments using Jalview, before being reported in a Microsoft^®^ Excel 2011 spread sheet and saved as a csv file. The phylogenetic trees shown alongside the cross-reactivity and the percentage identity tables were generated using Molecular Evolutionary Genetics Analysis (MEGA) software [22]: the aligned sequences were imported and trees derived using Unweighted Pair Group Method with Arithmetic mean (UPGMA), the simplest method of tree construction based on pairwise evolutionary distances. The trees generated were manually modified using MEGA and FigTree (http://tree.bio.ed.ac.uk/software/figtree/).

### Statistical analysis

Cross-reactivity tables for the IC_50_ neutralization titres, expressed as group, and for percentage amino acid identity, were completed using Microsoft^®^ Excel 2011. The R statistical software was then used to analyse the data and design a ‘ heat-map’ which colour codes the neutralization titres and the percentage identity. These codes are based on the use of the software package “RColorBrewer”, which permits building of a personalised colour palette, and “gplots”, a package that contains functions for the graphical interface. The “heatmap.2” function was eventually used to assign to each IC_50_ group or percentage identity value a colour. Kendall τ (tau) statistics (“Kendall” package) was also run using R software to check if association/correlation between measured IC_50_ titres and percentage identity was present.

